# *k*_inact_/*K*_I_ Value Determination for Penicillin-Binding Proteins in Live Cells

**DOI:** 10.1101/2024.05.05.592586

**Authors:** Joshua D. Shirley, Kelsie M. Nauta, Jacob R. Gillingham, Shivani Diwakar, Erin E. Carlson

**Author notes:** Co-authorship (Joshua D. Shirley, Kelsie M. Nauta).

## Abstract

Penicillin-binding proteins (PBPs) are an essential family of bacterial enzymes that are inhibited by the β-lactam class of antibiotics. PBP inhibition disrupts cell wall biosynthesis, which results in deficient growth and proliferation, and ultimately leads to lysis. IC_50_ values are often employed as descriptors of enzyme inhibition and inhibitor selectivity but can be misleading in the study of time-dependent, irreversible inhibitors. Due to this disconnect, the second order rate constant *k*_inact_/*K*_I_ is a more appropriate metric of covalent inhibitor potency. Despite being the gold standard measurement of potency, *k*_inact_/*K*_I_ values are typically obtained from *in vitro* assays, which limits assay throughput if investigating an enzyme family with multiple homologs (such as the PBPs). Therefore, we developed a whole-cell *k*_inact_/*K*_I_ assay to define inhibitor potency for the PBPs in *Streptococcus pneumoniae* using the fluorescent activity-based probe Bocillin-FL. Our results align with *in vitro k*_inact_/*K*_I_ data and show a comparable relationship to previously established IC_50_ values. These results support the validity of our *in vivo k*_inact_/*K*_I_ method as a means of obtaining a full picture of β-lactam potency for a suite of PBPs.

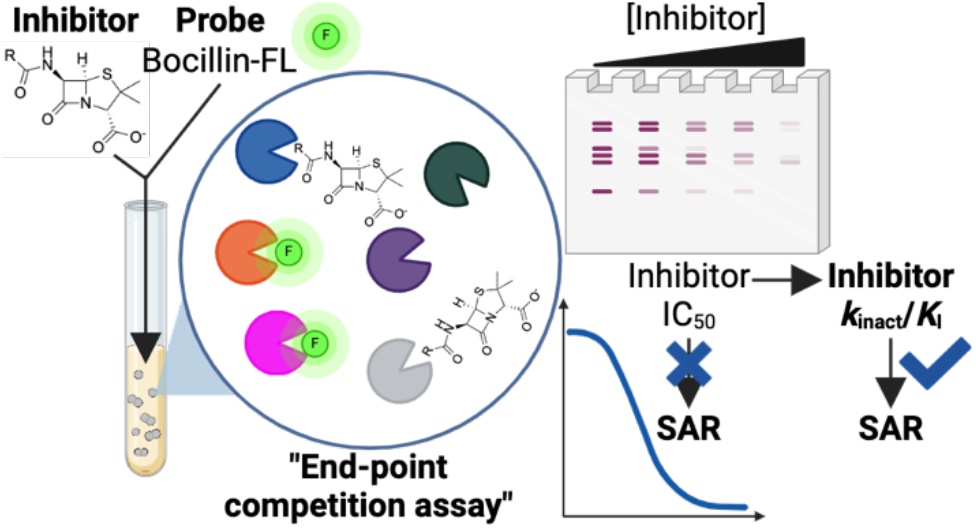

## Introduction

The penicillin-binding proteins (PBPs) are a family of membrane-associated bacterial enzymes inhibited by one of the most important and successful classes of antibiotics—the β-lactams.^1^ In bacteria, the PBPs catalyze three steps in cell wall biosynthesis: transglycosylase activity adds glycan monomers to growing chains of peptidoglycan (PG), transpeptidase activity crosslinks PG chains to form a 3-dimensional mesh, and peptidase activity modifies PG crosslink structure, as well as the availability of the PG pentapeptide for crosslinking.^2^ PBP inhibition by the β-lactams disrupts PG maintenance, leading to diminished cell proliferation and, ultimately, cell lysis.^3^

Inhibitor potency is commonly characterized by an IC_50_ value, or the inhibitor concentration required for 50% enzyme inhibition at incubation time *t*.^4-9^ Previous work in our lab has explored the use of β-lactams to define IC_50_ values for the PBP homologs in multiple Gram-positive and Gram-negative organisms.^10-14^ These studies aimed to identify inhibitors that would serve as scaffolds to develop activity-based probes (ABPs) selective for individual PBP homologs. While useful for our screen of relative PBP selectivity, the time and substrate concentration dependencies of IC_50_ values for irreversible inhibitors prohibit absolute inhibitor potency characterization for the study of structure-activity relationships (SAR).^9, 15, 16^ For example, with a sufficiently long incubation, most inhibitors will bind the target enzyme, making the resulting IC_50_ value a reflection of inhibitor concentration rather than inhibitor potency. Although strict standardization of experimental conditions allows for direct potency comparison using IC_50_ values, the second order rate constant *k*_inact_/*K*_I_ provides a time-independent potency measurement that can be used to conduct SAR studies for irreversible inhibitors (**Figure 1**).^17^

**Figure 1.**
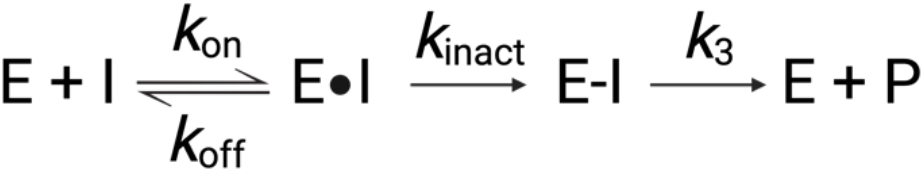
Enzymatic reaction steps for the binding of covalent inhibitors to the PBPs. *K*_I_ is dependent upon the rate of inhibitor association (*k*_on_), the rate of dissociation (*k*_off_) and *k*_inact_. *K*_I_, equal to (*k*_inact_ + *k*_off_)/*k*_on_, is distinct from the dissociation constant *K*_i_ (*k*_off_/*k*_on_).

The binding of irreversible inhibitors to the transpeptidase active site of the PBPs is initially driven by noncovalent interactions for the formation of the reversible Michaelis complex (E·I), in which the ligand moves in (*k*_on_) and out (*k*_off_) of the active site (**Figure 1**). The second step in the binding mechanism is acylation of the transpeptidase active site catalytic serine through nucleophilic attack of the electrophilic carbonyl group of the lactam ring; this step is defined by the first order rate constant *k*_inact_ (E-I). *K*_I_ is a metric of non-covalent binding affinity analogous to the Michaelis constant K_M_.^18-21^ Importantly, *K*_I_ is the concentration at which the pseudo-first order reaction rate *k*_obs_ is one-half of *k*_inact_, and is equal to (*k*_inact_+ *k*_off_)/*k*_on_.^22^ *K*_I_ should not be confused with *K*_i_, which is the dissociation constant for an irreversible inhibitor (equal to *k*_off_/*k*_on_ and analogous to *K*_d_).^22^ Despite their different definitions, both *K*_I_ and *K*_i_ are commonly used to characterize inhibitor potency.^22, 23^ Finally, the deacylation constant, or hydrolysis rate, is defined by *k*_3_, when the acylated intermediate is hydrolyzed and the catalytic serine is restored. For “irreversible” PBP inhibitors, *k*_3_ is often negligible, as the hydrolysis rates are slow, with values for β-lactam-susceptible PBPs in the range of 10^-6^ s^-1^.^24-26^

Determination of inhibitor *K*_I_ and *k*_inact_ values is the most descriptive characterization of the contributions of the non-covalent and covalent binding steps to the potency of PBP inhibitors. The ratio *k*_inact_/*K*_I_ (analogous to the catalytic efficiency quotient *k*_cat_*/K*_M_) is the second-order rate constant of inhibition regarded as the gold standard for time-dependent, irreversible inhibitor potencies.^18, 19, 27^ Whether through calculation of *k*_inact_ and *K*_I_ values or through direct determination of *k*_inact_*/K*_I_, current methods typically use recombinantly expressed and purified proteins.^26, 28-40^ This *in vitro* approach is limited in both its throughput and by its lack of biological context (i.e., *in vitro* studies often use recombinant, truncated PBPs and may not reflect the confirmation found *in vivo*). Therefore, the development of a live-cell *k*_inact_/*K*_I_ assay is crucial for accurately characterizing β-lactam inhibitor potency in the context of a SAR campaign.^18, 19^

To date, *in vitro k*_inact_/*K*_I_ experiments with PBPs have proven useful for the screening and development of inhibitors against the PBPs of interest; however, there has been only one report of an assay capable of measuring *k*_inact_/*K*_I_ values for an inhibitor against the entire PBP profile of an organism.^41^ This work obtained *k*_inact_/*K*_I_ values for the PBPs of penicillin-susceptible and -resistant *Staphylococcus aureus* strains through incubation of membrane fractions with increasing [^3^H]-penicillin concentrations. While this study clearly represents an important step closer to investigations performed *in vivo*, our work and others has demonstrated significant differences in enzyme function and conformation following cell lysis.^42, 43^ In addition, while most *in vitro* assays use fluorescence or mass spectrometry as a readout, this work used a radiolabel and also relied on time-consuming classical enzyme kinetics strategies, requiring multiple inhibitor concentrations and incubation times.^26, 41, 44, 45^

To efficiently measure potency values for the entire PBP profile of an organism, improved throughout is required. Here, we report the development of a gel-based, end-point competition assay that is performed on live cells and enables rapid determination of *k*_inact_/*K*_I_ values for all enzymatically active PBP homologs. Our approach was informed by a previously described assay reported by Miyahisa *et. al*. to determine *k*_inact_/*K*_I_ values for inhibitors of epidermal growth factor receptor (EGFR).^46^ We calculated *k*_inact_/*K*_I_ values for ten β-lactam antibiotics for the native PBPs of *Streptococcus pneumoniae* IU1945 (D39 /Δ*cps*);^47^ these results aligned with previously reported *in vitro* potency values.^26^ Additionally, correlation of β-lactam *k*_inact_/*K*_I_ values and our reported live cell IC_50_ values from identical incubation times validates our approach as a powerful tool for PBP characterization.^10, 13^

## Results and Discussion

### Determination of *k*_inact_/*K*_I_ values for Bocillin-FL against PBPs in *S. pneumoniae*

To perform the required competition assay, we first needed to determine the *k*_inact_/*K*_I_ value for a universal probe, such as the fluorescent penicillin, Bocillin-FL, against each PBP.^48^ This probe would then serve as a competitor against other β-lactams in a time-course fluorescence experiment on live cells – simultaneously enabling the investigation of all six PBP homologs in *S. pneumoniae*. To obtain the Bocillin-FL *k*_inact_/*K*_I_ values, this probe was added to *S. pneumoniae* cells in PBS and aliquots of the reaction mixture were removed at respective timepoints and quenched in acetonitrile with 1% formic acid, conditions adapted from a stopped-flow method utilized by Lu *et. al*.^26^ Following cell lysis and protein separation by SDS-PAGE, we observed time-dependent increases in fluorescence, suggesting that this quenching method was sufficient to stop enzymatic activity (**Figure 2A**). We calculated *k*_obs_ by measuring the fluorescence of increasing concentrations of Bocillin-FL over time in whole cells and using Equation 1 (**Figure 2B**).^24^

**Figure 2.**
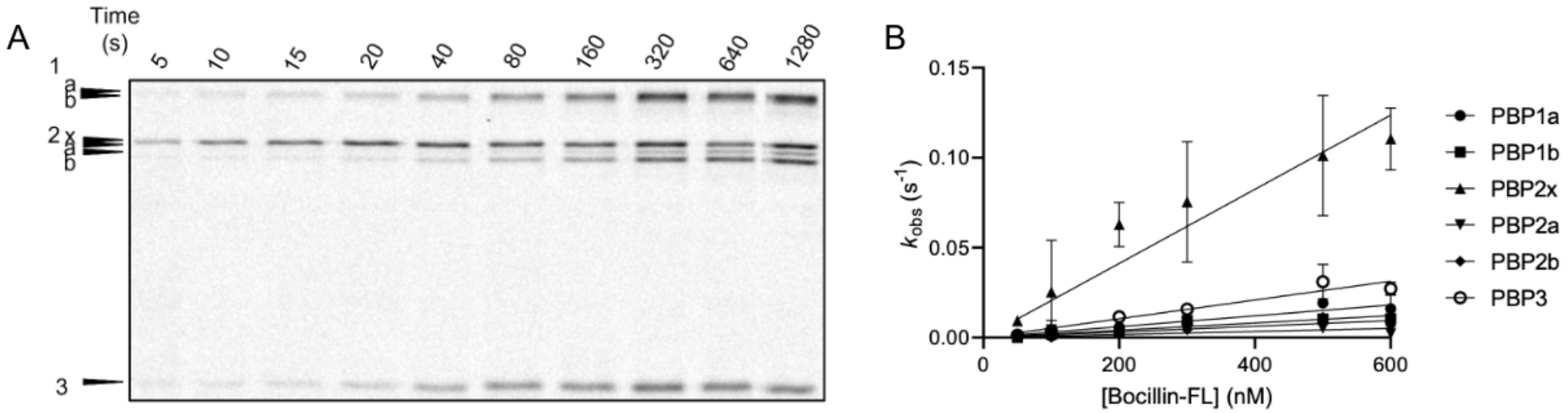
**A**. Time-dependent inhibition of PBPs in live *S. pneumoniae*. Live cells were treated with 300 nM Bocillin-FL and at respective timepoints, aliquots of the reaction were removed and quenched in acetonitrile with 1% formic acid. SDS-PAGE analysis revealed that this is an adequate quenching method and reaction rates can be determined. **B**. Linear regression analysis of *k*_obs_ vs Bocillin-FL for PBPs in live *S. pneumoniae*. Live cells were treated with increasing concentrations of Bocillin-FL and *k*_obs_ rate constants were calculated from respective progress curves. Mean and standard deviation for *k*_obs_ were calculated from duplicate experiments.

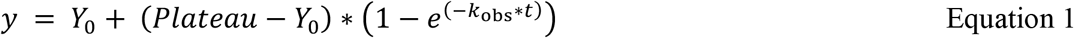

Where *y* is the fluorescence intensity at a given time, *t, Y*_0_ is the fluorescence intensity at zero minutes, *plateau* is the fluorescence intensity at enzyme saturation, and *k*_obs_ is the pseudo-first order rate constant.

At low inhibitor concentration (i.e., orders of magnitude lower than *K*_i_), the relationship between *k*_obs_ and inhibitor concentration will be linear, with the slope equal to *k*_inact_/*K*_I_.^18, 27^ For facile *k*_inact_/*K*_I_ determination, we used Bocillin-FL concentrations in the nanomolar range, which is well below the reported millimolar *K*_i_ values for penicillin G with *S. pneumoniae* PBP2x (0.9 mM and 20 mM from separate studies).^26, 33, 49^ As such, we cannot accurately determine the individual *k*_inact_ and *K*_I_ values, but we can determine the ratio *k*_inact_*/K*_I_ using linear least squares regression analysis of the *k*_obs_ vs [Bocillin-FL] plot (**Table 1**).^27^ Comparing reported *k*_inact_*/K*_I_ values of β-lactams determined using pure PBPs, we demonstrated good agreement with our calculated *k*_inact_/*K*_I_ values for each of the six *S. pneumoniae* PBP homologs. For example, our calculated Bocillin-FL *k*_inact_/*K*_I_ for PBP2x (206,000 ± 29,800 M^-1^s^-1^) is comparable to a *k*_inact_/*K*_i_ value reported for the closely related penicillin G (200,000 M^-1^s^-1^).^26^ These results demonstrate that our live cell kinetics assay can produce reliable *k*_inact_/*K*_I_ values for Bocillin-FL against the entire PBP profile in *S. pneumoniae*.

**Table 1.** *k*_inact_/*K*_I_ values obtained from live cell Bocillin-FL kinetics assays.

| PBP | $k_{\text{inact}}/K_I$ ($\text{M}^{-1}\text{s}^{-1}$)<sup>a</sup> | Literature β-Lactam<br>$k_{\text{inact}}/K_I$ ($\text{M}^{-1}\text{s}^{-1}$)<sup>b</sup> | Reference |
| --- | --- | --- | --- |
| PBP1a | $30,200 \pm 5,800$ | 34,200 - 69,000 | 28, 29 |
| PBP1b | $20,300 \pm 1,200$ | 19,509 - 20,431 | 30 |
| PBP2x | $206,000 \pm 30,000$ | 58,000 - 200,000 | 26, 31-34 |
| PBP2a | $8,500 \pm 3,500$ | 17 - < 3,800 | 26, 35, 36 |
| PBP2b | $15,700 \pm 500$ | 930 - 13,700 | 37 |
| PBP3 | $52,300 \pm 9,400$ | 10,300 - 30,000<sup>c</sup> | 38, 39 |

<sup>a</sup>Average and standard deviations from biological duplicates
<sup>b</sup>Literature values were determined *in vitro* with penicillin G and/or Bocillin-FL
<sup>c</sup>Values were reported for DD-carboxypeptidases from species other than *S. pneumoniae*

### Bocillin End-Point Competition Assay

Following determination of the *k*_inact_/*K*_I_ parameter of Bocillin-FL for respective PBPs in *S. pneumoniae*, we determined the potency of β-lactams against the entire PBP complement using our Bocillin-FL *k*_inact_/*K*_I_ values in combination with β-lactam IC_50_ values obtained from a gel-based, end-point competition assay. Live cells were simultaneously treated with increasing concentrations of inhibitor and Bocillin-FL. The end-point time was defined as the amount of time it takes to occupy 99% of a given PBP (*t*_99_), which was calculated from the *k*_obs_ values using Equation 4 (derived from Equations 2 and 3).

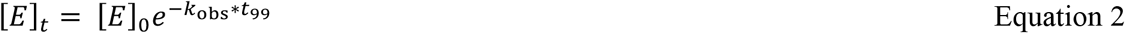

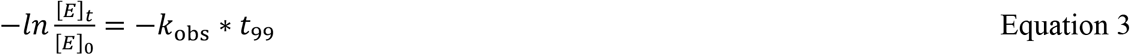

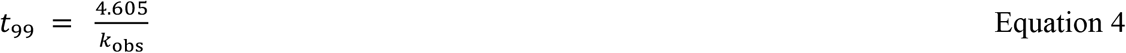

Where *t*_99_ represents the time at which 99% of the enzyme is occupied, *[E]*_*t*_ is the percent unoccupied enzyme at *t*_*99*_ (1%) and *[E]*_*0*_ is the percent of unoccupied enzyme at *t*_0_ (100%), and *k*_obs_ is the pseudo-first order rate constant calculated for the specific concentration of Bocillin-FL.^18, 40^ Due to the slow hydrolysis of the acyl-enzyme complex (*k*_3_ ∼ 10^-6^ s^-1^), we used an end-point time that was derived from the slowest *k*_obs_ value obtained from the six PBPs, which was PBP2a 0.0022 s^-1^ for 600 nM Bocillin-FL.^26^ In doing so, all six PBPs will have 99% active site occupancy, mitigating the need to run experiments using different end-points for each PBP. IC_50_ values could then be calculated from these data, and the value of inhibitor *k*_inact_/*K*_I_ determined from Equation 5.^46^

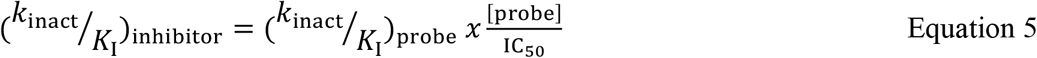

To do this, we first generated IC_50_ values for each PBP homolog in *S. pneumoniae* with a given inhibitor. Briefly, live cells were incubated with 600 nM Bocillin-FL and increasing concentrations of respective inhibitor (10^-4^-10^4^ µM) for 30 min. Following cell lysis and centrifugation, the membrane proteome was resolved on an SDS-PAGE gel; IC_50_ values were calculated as previously described (**Figures 3, S1-S11**).^10^ As in previous work, strains lacking either PBP1a (*Δpbp1a*) or PBP1b (*Δpbp1b*) were included to enable accurate quantitation of labeled proteins in this region of the gel (**Figure 3B** and **C**).^10^ Ten β-lactams in total were tested using this method (**Table 2**). Our panel includes 7 penicillins and 3 cephalosporins, encompassing a wide range of *S. pneumoniae* MIC values (0.08-4 µg/mL), IC_50_ values (most potent PBP inhibition 7.2 nM-220 µM), and PBP isoform selectivities.^10, 13^

**Table 2.** Live-cell *k*_inact_/*K*_I_ values for β-lactam inhibition of the *S. pneumoniae* PBPs.

| Inhibitor | $k_{inact}/K_i$ (M<sup>-1</sup> s<sup>-1</sup>) (Mean ± SD) | | | | | | PBP selectivity by IC<sub>50</sub><sup>10, 13</sup> |
| --- | --- | --- | --- | --- | --- | --- | --- |
|  | PBP1a | PBP1b | PBP2x | PBP2a | PBP2b | PBP3 |  |
| Penicillin |  |  |  |  |  |  |  |
| Penicillin G | 70,000 ± 20,000 | 41,000 ± 11,000 | 200,000 ± 7,400 | 15,000 ± 5,400 | 11,000 ± 2,300 | 180,000 ± 55,000 | 2x, 3 |
| Ampicillin | 15,000 ± 1,300 | 2,000 ± 310 | 59,000 ± 6,600 | 4,600 ± 480 | 6,700 ± 240 | 77,000 ± 14,000 | 1a, 2x, 3 |
| Dicloxacillin | 12,000 ± 990 | 2,800 ± 780 | 38,000 ± 5,200 | 2,000 ± 100 | 3,300 ± 620 | 32,000 ± 5,100 | 2x, 3 |
| Mecillinam | 18 ± 0.50 | 56 ± 18 | 410 ± 58 | 62 ± 2.7 | 26 ± 1.8 | 620 ± 150 | 1a, 1b |
| Methicillin | 5.4 ± 1.2 | 650 ± 430 | 26,000 ± 4,900 | 160 ± 21 | 780 ± 65 | 4,800 ± 520 | 2x, 3 |
| Piperacillin | 1,800 ± 190 | 2,500 ± 620 | 140,000 ± 48,000 | 880 ± 75 | 15,000 ± 810 | 24,000 ± 3,500 | 2x, 3 |
| <u>Cephalosporin</u> |  |  |  |  |  |  |  |
| Cephalexin | 220 ± 120 | 220 ± 83 | 360 ± 210 | 81 ± 12 | 4 ± 2.7 | 3,700 ± 980 | 3 |
| Cefotaxime | 10,000 ± 5,000 | 52,000 ± 4,000 | 730,000 ± 58,000 | 16,000 ± 4,700 | 5.4 ± 1.3 | 1,400,000 ± 460,000 | 2x, 3 |
| Cefsulodin | 85 ± 6.5 | 170,00 ± 2,400 | 1,000 ± 6 | 370 ± 7 | >RT | 3,300 ± 250 | 3 |
| Ceftriaxone | 1,700 ± 170 | 9,200 ± 3,200 | 30,000 ± 7,800 | 610 ± 280 | >RT | 3,700 ± 980 | 2x, 3 |

RT = Range tested

**Figure 3.**
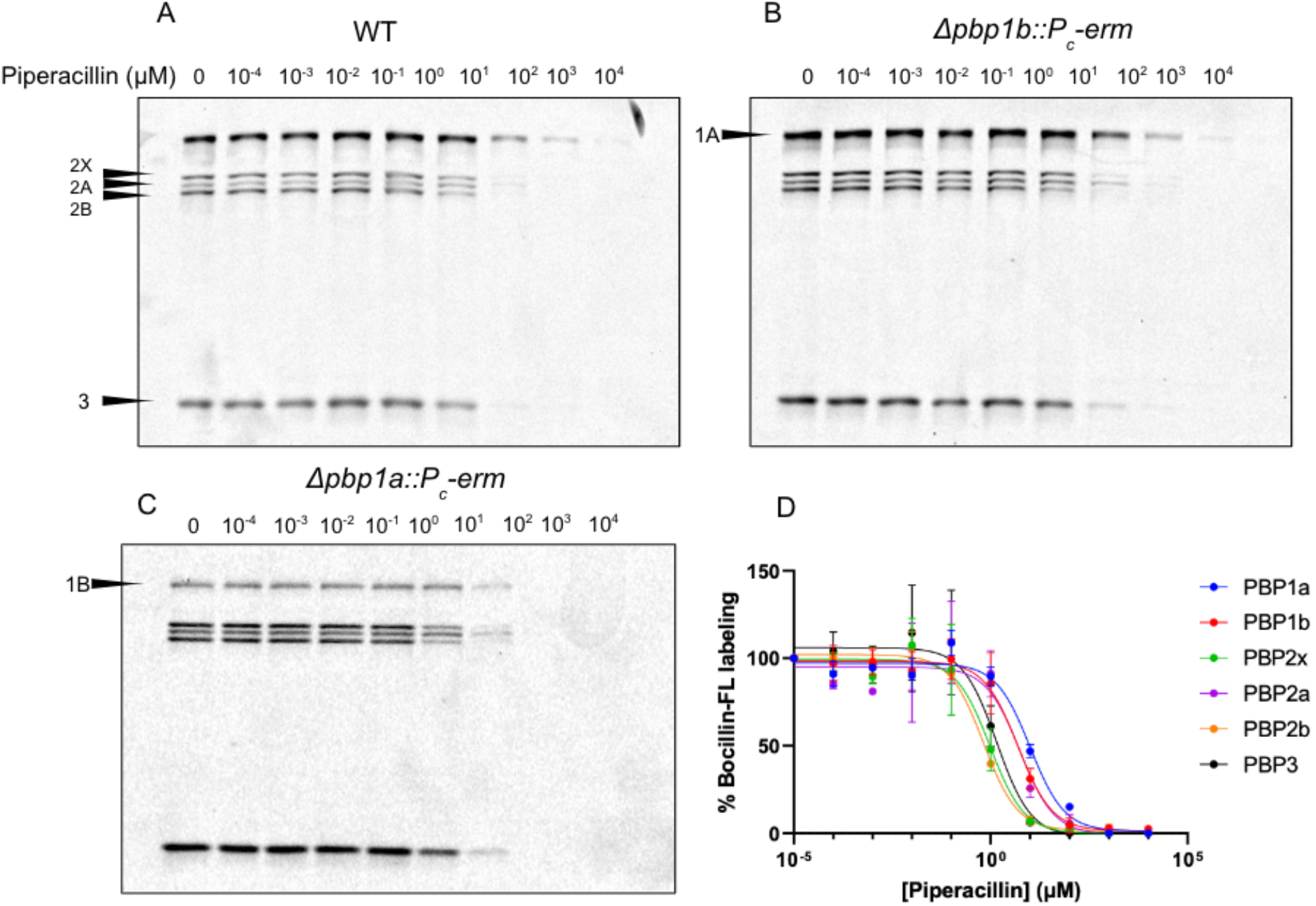
Bocillin end point assay representative gel images. Live *S. pneumoniae* WT (**A**), *Δpbp1b* (**B**), or *Δpbp1a* (**C**) cells were treated with Bocillin-FL (600 nM) and a concentration gradient of piperacillin (10^-4^-10^4^ µM). After incubation, cells were lysed and isolated membrane fractions were run on an SDS-PAGE gel. **D**. Quantification of dose-dependent inhibition of PBPs. The data were fit to a nonlinear regression (curve fit) using log[inhibitor] vs. response (three parameter) in GraphPad Prism. Contrast and brightness were optimized using ImageJ uniformly over each gel and IC_50_ values were calculated using GraphPad Prism.

To assess the relationship between *k*_inact_/*K*_I_ and IC_50_ values as metrics of inhibitor potency, we plotted the *k*_inact_/*K*_I_ values for each PBP homolog against our published IC_50_ values (**Figure 4**). Previously, Maurer *et al*. derived a time-and substrate concentration-dependent algebraic model connecting IC_50_ and *k*_inact_/*K*_i_.^15^ The relationship between values of IC_50_ and *k*_inact_/*K*_i_ is inverse as a high *k*_inact_/*K*_i_ corresponds to high potency, while a high IC_50_ corresponds to low inhibition. Subsequent investigations of Janus kinase, kynurenine aminotransferase II, and KRAS^G12C^ covalent inhibitors support this model.^17, 23, 50, 51^ As *K*_I_ = *K*_i_ + (*k*_inact_/*k*_on_), we reasoned that this relationship would extend to *k*_inact_/*K*_I_.^22^ Accordingly, live cell data shows positive associations between pIC_50_ and log(*k*_inact_/*K*_I_), which corresponds to an inverse correlation between IC_50_ and *k*_inact_/*K*_I_ (**Figure 4**). Furthermore, we observed only minor trend differences in inhibitor selectivity using *k*_inact_/*K*_I_ values compared to selectivity determination by IC_50_. While IC_50_ and *k*_inact_/*K*_I_ values concur fairly well in comparisons of covalent inhibition, the dependence of IC_50_ values on incubation time limits their utility to future SAR campaigns. Therefore, the devised fluorescent gel-based assay for live cell PBP *k*_inact_/*K*_I_ value determination has great potential.

**Figure 4.**
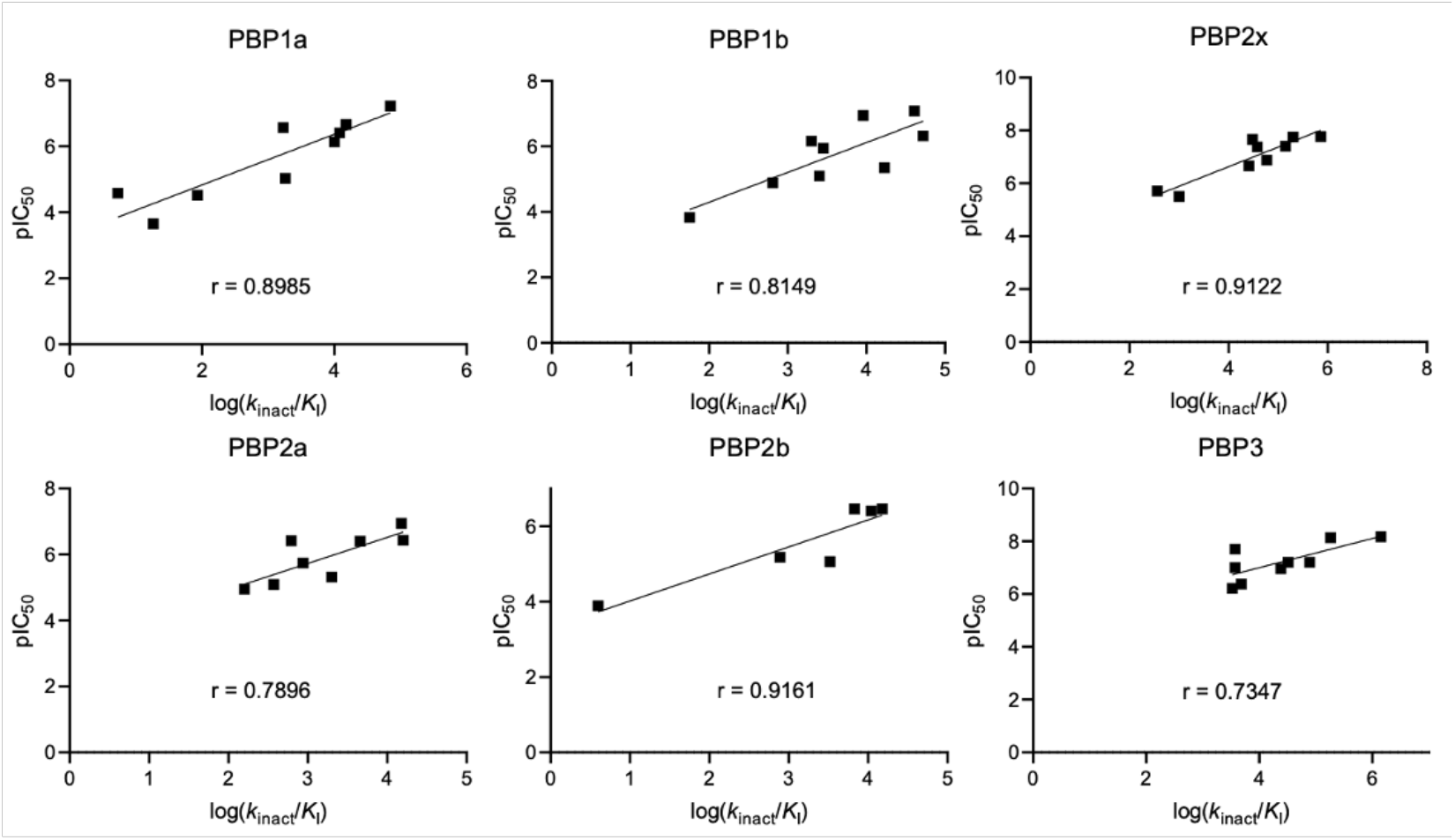
Correlation analysis of live-cell log(*k*_inact_/*K*_I_) and pIC_50_ values for PBP inhibition in *S. pneumoniae*. Inhibition metrics for the ten β-lactams assessed with both assays are plotted for each PBP isoform.

We propose that the distribution of β-lactam potency arises from variations in *K*_I,_meaning that the chemical diversity of β-lactam substituents substantially affects Michaelis complex formation. This model is supported by a high-throughput screen of covalent inhibitors of the L,D-transpeptidase Ldt_Mt2_ in *Mycobacterium tuberculosis*.^52^ Here, it was shown that the intrinsic reactivity of the warhead (*k*_chem_) remained relatively static when plotted against *k*_inact_/*K*_I_, implying that noncovalent association drives changes to covalent inhibitor potency. Furthermore, Lu and colleagues demonstrated similar *k*_inact_ values for PBP2x inhibition with penicillin G and cefotaxime (180 s^-1^ and 150 s^-1^, respectively), suggesting that the rapid kinetics of the acylation step are not profoundly affected by inhibitor structure.^26^ This work demonstrates that variations in covalent inhibitor potency are likely mediated by noncovalent interactions, as opposed to the electrophilicity of the covalent warhead.

To evaluate the assay’s utility for future SAR campaigns, we sought to compare predictions of *K*_I_ from our method to noncovalent affinity values previously determined from *in vitro* experiments. This was achieved with published *k*_inact_ values using Equation 6.

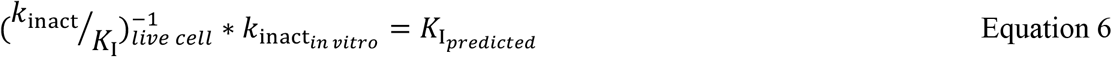

We used delineated *k*_inact_ and *K*_i_ values in *S. pneumoniae* from Lu *et al*.^26^ Live cell *k*_inact_*/K*_I_ values aligned with their *in vitro* results for PBP2x; for example, Equation 6 generated a *K*_I_ of 0.9 mM for penicillin G, identical to the reported *K*_i_. Our predicted *K*_I_ for PBP2x cefotaxime inhibition also agreed with the results of this study. The predicted *K*_I_ for penicillin G inhibition of PBP2a (150 µM) shows a difference of several orders of magnitudes compared to the *in vitro K*_i_ value (13 mM). This discrepancy is likely due either to the distinct definitions of *K*_I_ and *K*_i_, or *in vivo* modulation of enzyme activity and active site accessibility. Activity regulation is particularly likely as PBP2a is a bifunctional transglycosylase/transpeptidase.^53, 54^ While our live cell assay provides the most realistic view of inhibition in the biological context, alignment with *in vitro* potency measurements serves as a key validation step.

## Conclusion

The data presented here support the validity of our novel *in vivo* assay in *S. pneumoniae* for defining *k*_inact_*/K*_I_ of a universal fluorescent probe for PBPs, as well as a small library of β-lactam inhibitors. Comparison of our results to previously reported values of *in vitro k*_inact_*/K*_I_ values for β-lactams binding to PBPs of *S. pneumoniae* reveals good agreement with our cell-based assay. This agreement provides validation for the use of the live cell assay, which will enable SAR studies in a physiologically relevant environment and mitigate the need to produce an entire profile of recombinant PBPs for any given organism (which can be as many as 18 PBP homologs). We also observed a relationship between our previously published IC_50_ values in *S. pneumoniae* and the *k*_inact_/*K*_I_ values presented here. IC_50_ values are easily obtained and offer valuable insight into selectivity; however, they are insufficient descriptors of potency due to their time dependency and inability to delineate Michaelis complex formation and acylation. Therefore, application of this or a similar assay to determine *k*_inact_/*K*_I_ values must be considered to properly inform SAR campaigns and the design of time-dependent irreversible inhibitors.

## Materials and Methods

### Bacterial Strains, Growth Conditions, and Antibiotics

*S. pneumoniae* strain IU1945 (a derivative of D39 (*Δcps*)),^47^ E177 (*Δpbp1a::P*_*c*_*-erm*), and E193 (*Δpbp1b::P*_*c*_*-erm*) (gifts from Malcolm E. Winkler, Department of Biology, Indiana University-Bloomington, USA) were incubated statically in BBL Brain Heart Infusion Broth (Becton, Dickinson, and Company, Sparks, MD 21152 USA, REF 211059) at 37 °C overnight in 5% CO_2_.^55^ Cultures at an OD_620_ of 0.15-0.20 were subcultured 1:10 into fresh BHI and incubated to an OD_620_ of 0.25-0.3.

Cefsulodin, dicloxacillin, mecillinam, methicillin, penicillin G, and piperacillin were purchased from Sigma Aldrich (St. Louis, MO). Ceftriaxone and cephalexin were purchased from Research Products International (Mt. Prospect, IL). Ampicillin and cefotaxime were purchased from Calbiochem (Billerica, MA). Bocillin-FL was purchased from Invitrogen (Waltham, MA). All antibiotics were stored as solids at the temperature indicated by the manufacturer and were dissolved in 1x PBS at 0.04 M. Once dissolved, antibiotics were stored at -20 °C. All stock solutions were diluted 1:2 with 2400 nM Bocillin-FL in PBS to make 2000 µM antibiotic, 1200 nM Boc-FL stocks. They were then serially diluted 1:10 in 1200 nM Bocillin-FL to make 0.0002-2,000 µM working solutions.

### Whole-cell Bocillin-FL Time Course Assays

Fresh cultures of *S. pneumoniae* were grown as described above. Once cultures reached an OD_620_ = 0.2 – 0.3, 1 mL was added to a 1.7 mL Eppendorf tubes and was harvested by centrifugation (16,000 x *g*, 2 min, RT). Cell pellets were washed with 1 mL 1X PBS (pH 7.4) and collected by centrifugation. A total of 6 mL PBS was used to resuspend the collective cell pellets, resulting in 6 mL of resuspended cells in PBS with an OD_620_ = ∼0.3. To this suspension was added respective amounts of 5 mM Bocillin-FL (100% DMSO) to afford the desired final concentration. At each time point (5, 10, 15, 20, 40, 80, 160, 320, 640, 1280 s), 500 µL of the reaction was aliquoted into respective Eppendorf tubes containing 200 µL 99% acetonitrile/1% formic acid. Once all time points had been collected, samples were centrifuged at 21,000 x *g* for 10 min. The supernatant was removed and discarded. Pellets were resuspended in 100 µL 50 mM Tris pH 7.8 – 10 mg/mL lysozyme and incubated at 37 °C for 20 min. Samples were lysed on a vial tweeter (90% C, 95% A, 5% adjustment snap, 50 s on, 10 s off for 6 min total) and the membrane fraction was collected by centrifugation at 21,000 x *g* for 10 min at 4 °C. The supernatant was removed and discarded, and the membrane fractions were resuspended in 34 µL 50 mM Tris pH 7.8 – 0.5% SDS. Protein measurements were taken on by UV-Vis using a NanoDrop and recorded. Ten microliters of 4X SDS loading buffer (200 mM Tris base, 400 mM DTT, 8% SDS, 0.4% Bromophenol Blue, 40% glycerol, pH 6.8) were added to each tube and the samples were boiled at 95 °C for 5 min. Samples were cooled and vortexed. Based on the protein concentration of each sample, 80 µg of protein was loaded onto the gel (volume added depended on the protein concentration). Gels were run for 1 ½ hours, or until the SDS loading buffer had run off the foot of the gel (180 V, 60 W, 40 mA/gel).

Gels were removed from their cassettes, rinsed twice with Milli-Q water, and analyzed using a GE Typhoon FLA 9500 gel scanner using the BODIPY-FL settings. Integrated density measurements were taken for fluorescence and Coomassie stains. The normalized fluorescence intensities were added into an XY table in GraphPad Prism and *k*_obs_ values were calculated using Equation 1. Each *k*_obs_ and corresponding Bocillin-FL concentration for each PBP were entered into an XY plot in GraphPad prism and analyzed using linear regression. The slope of the line for each PBP was determined and used as the *k*_inact_/*K*_I_ value for Bocillin-FL against each of the six PBPs in *S. pneumoniae*.

### Bocillin-FL Competition End Point Assay

Cells grown to an OD_620_ of 0.25-0.30 were centrifugated at 1,503 x *g* for 10 min and washed once in PBS. The cells were pelleted and incubated in PBS containing Bocillin-FL (600 nM) and β-lactam inhibitors at 10^4^-10^-4^ µM for 30 min at RT. The cells were incubated with lysozyme (10 mg/mL) in 50 mM TRIS for 15 minutes at 37 °C. The cells were lysed using a Hielscher ultrasonic processor UP200St (90% C, 95% amplitude, 5% adjustment snap, 50 s on, 10 s off, for 6 min on ice). The membrane proteome was isolated by centrifugation at 21,000 x *g* for 10 min at 4 °C. The pellets were resuspended in 0.5% SDS in 50 mM TRIS. The protein concentrations were measured using a NanoDrop 1000 spectrophotometer (Thermo Scientific, Wilmington, DE). SDS loading buffer was added, and the proteins were incubated at 90 °C for 3-5 min. Protein (80 µg) was run on a 10% SDS-PAGE gel. The protein bands were separated by gel electrophoresis for ∼2.5 h at constant 180 V, 400 mA (maximum), and 60 W. The gels were rinsed with DI water and imaged using a Typhoon 9500 gel scanner (Amersham Biosciences, Pittsburgh, PA) with a 473 nm laser excitation with LPB filter (≥510 nm) and imaged at 50 μm resolution. After imaging, the gels were stained with Coomassie blue (BioRad), heated for 30 s in a microwave, and incubated at RT for 30 min. The gels were transferred to destain (10% acetic acid, 40% methanol, 50% H_2_O and heated in a microwave for 30 s. The gels were incubated overnight and imaged the next day using a Typhoon 9500 gel scanner with a 635 nm laser excitation with LPR filter (≥665 nm) at a 50 μm resolution.

### Image Processing

The gels were analyzed using ImageJ 1.53k (National Institute of Health, MD). The background was subtracted, and the signal-to-noise ratio was optimized by adjusting the brightness and contrast of the entire gel. The integrated density was measured to quantify protein band intensity. The integrated densities were normalized to total protein integrated densities measured from 4 protein bands from Coomassie blue gels. Percent of the untreated control was calculated. GraphPad Prism 9.4.1 (681) (La Jolla, CA) was used to calculate the IC_50_ values and fit the data to a nonlinear regression (curve fit) using log(inhibitor) vs. response (three parameter). *k*_inact_/*K*_I_ values and Pearson correlation coefficients were calculated in GraphPad Prism. All graphs were made in GraphPad Prism.

## Author information

J.D.S. conceived of and designed this project, generated data for Figure 2, wrote a portion of the manuscript draft, and edited the manuscript. K.M.N. generated data for Figure 3, Table 2, and Figures S1-S11, performed data analysis for Figure 4, wrote a portion of the manuscript draft, and edited the manuscript. J.R.G. wrote a portion of the manuscript draft and performed data analysis for Figure 4. S.D. assisted with data generation for Figure 3 and Table 2. E.E.C. contributed to project design, data analysis, and manuscript editing.

The authors declare no competing financial interest.

## Acknowledgements

This work was supported by the National Institutes of Health (R01 GM128439 A1, E.E.C.) and the University of Minnesota, Department of Chemistry. J.D.S. was supported by the National Institutes of Health’s National Center for Advancing Translational Sciences, grants TL1R002493 and UL1TR002494. The content is solely the responsibility of the authors and does not necessarily represent the official views of the National Institutes of Health’s National Center for Advancing Translation Sciences. J.R.G. was supported by the National Institutes of Health/National Institute of General Medical Sciences Chemistry Biology Interface Training Grant 5T32GM132029-5. Graphical figures created with GraphPad Prism and BioRender.

