## Supporting Information Document for "*k*_inact_/*K*_I_ Value Determination for Penicillin-Binding Proteins in Live Cells"

[f]*Co-authorship*

**Corresponding Author*

**Table of contents**

**Figure S1.** Quantification of dose-dependent inhibition of PBPs S2

**Figure S2.** Bocillin End Point Assay representative gel images for ampicillin inhibition S3

**Figure S3.** Bocillin End Point Assay representative gel images for methicillin inhibition S4

**Figure S4.** Bocillin End Point Assay representative gel images for dicloxacillin inhibition S5

**Figure S5.** Bocillin End Point Assay representative gel images for penicillin G inhibition S6

**Figure S6.** Bocillin End Point Assay representative gel images for cefsulodin inhibition S7

**Figure S7.** Bocillin End Point Assay representative gel images for cephalexin inhibition S8

**Figure S8.** Bocillin End Point Assay representative gel images for ceftriaxone inhibition S9

**Figure S9.** Bocillin End Point Assay representative gel images for piperacillin inhibition S10

**Figure S10.** Bocillin End Point Assay representative gel images for cefotaxime inhibition S11

**Figure S11.** Bocillin End Point Assay representative gel images for mecillinam inhibition S12

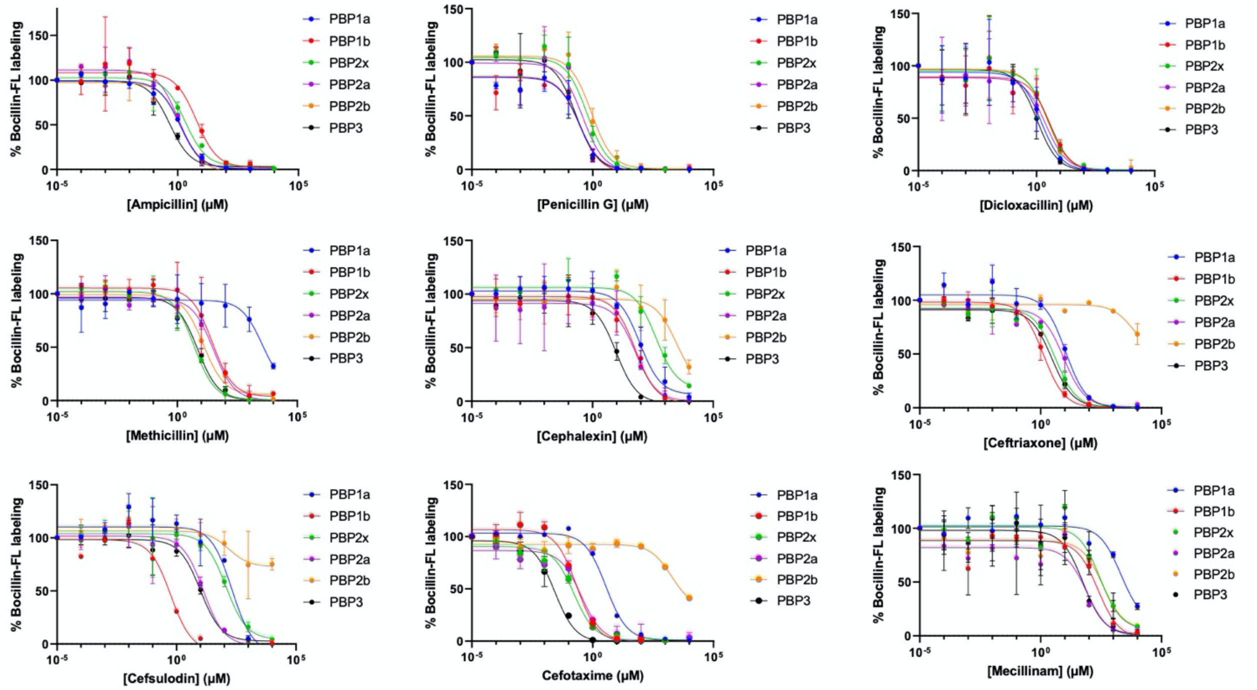


**Figure S1.** Quantification of dose-dependent inhibition of PBPs. This figure includes all β-lactams tested in this study. The data were fit to a nonlinear regression (curve fit) using log(inhibitor) vs. response (three parameter) in GraphPad Prism. Contrast and brightness were optimized using ImageJ and $IC_{50}$s were calculated using GraphPad Prism.

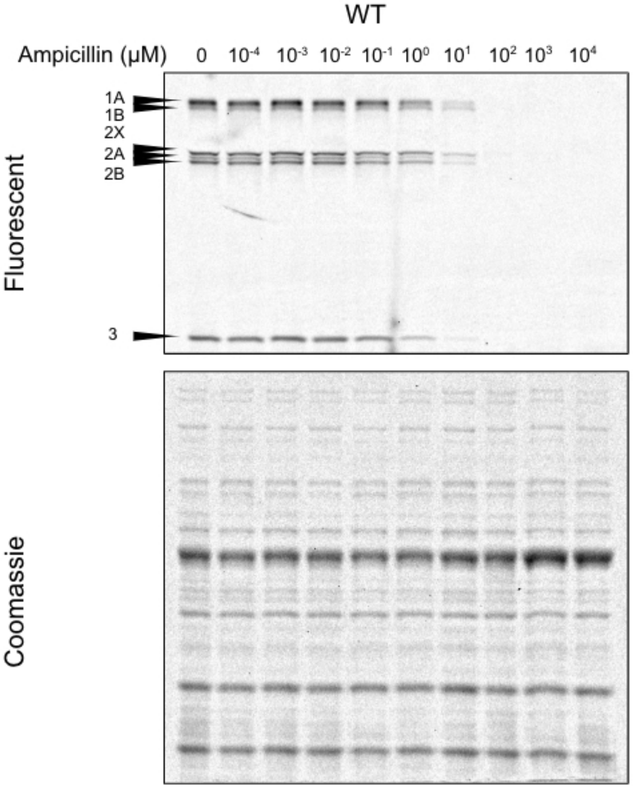


**Figure S2**. Bocillin End Point Assay representative gel images for ampicillin inhibition. Live *S. pneumoniae* WT cells were treated with Bocillin-FL (600 nM) and a concentration gradient of ampicillin ($10^{-4}$-$10^{4}$ μM). After incubation, the cells were lysed with lysozyme and sonication. The membrane fractions were isolated and 80 μg per sample were run on an SDS-PAGE gel and imaged for Bocillin-FL (top). After imaging the gels were Coomassie stained for total protein quantification (bottom). See Figure S1 for quantification.

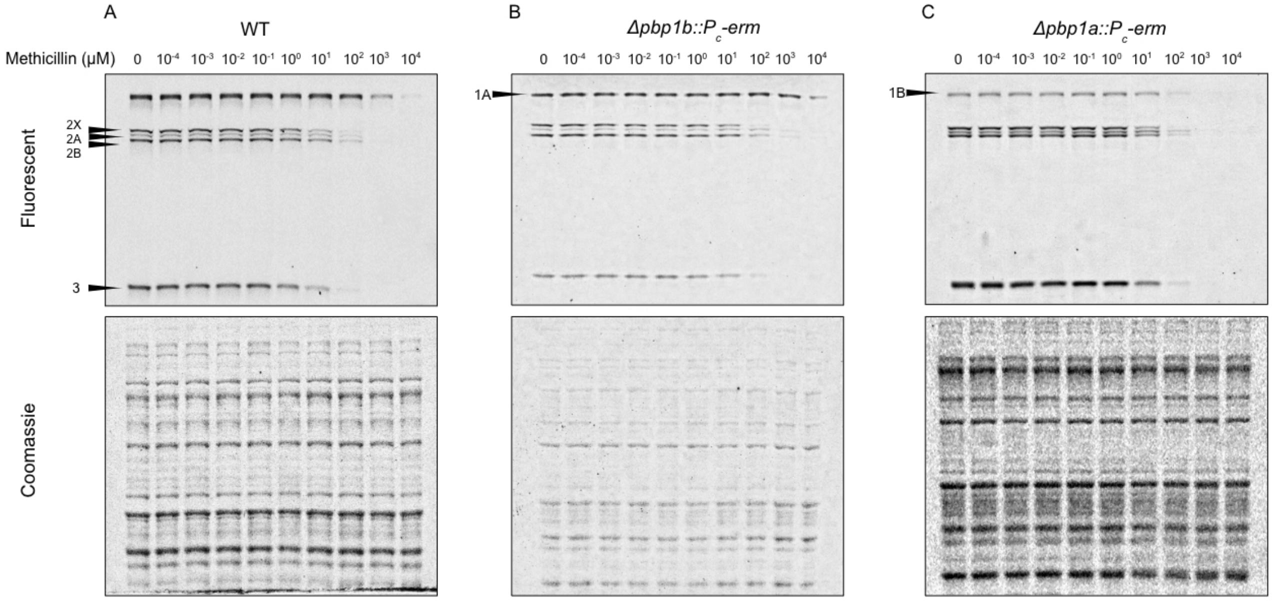


**Figure S3**. Bocillin End Point Assay representative gel images for methicillin inhibition. Live *S. pneumoniae* WT, Δ*pbp1B* (B), or *Δpbp1A* (C), cells were treated with Bocillin-FL (600 nM) and a concentration gradient of methicillin ($10^{-4}$-$10^{4}$ μM). After incubation, the cells were lysed with lysozyme and sonication. The membrane fractions were isolated and 80 μg per sample were run on an SDS-PAGE gel and imaged for Bocillin-FL (top). After imaging the gels were Coomassie stained for total protein quantification (bottom). See Figure S1 for quantification.

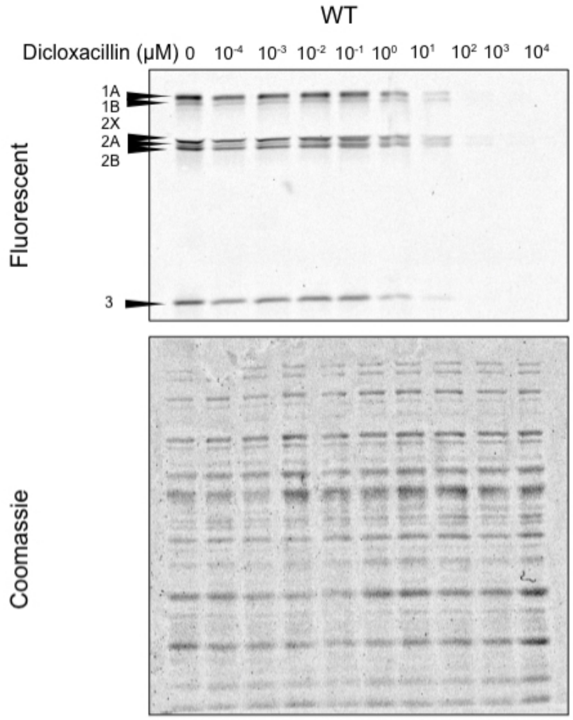


**Figure S4**. Bocillin End Point Assay representative gel images for dicloxacillin inhibition. Live *S. pneumoniae* WT cells were treated with Bocillin-FL (600 nM) and a concentration gradient of dicloxacillin ($10^{-4}$-$10^{4}$ μM). After incubation, the cells were lysed with lysozyme and sonication. The membrane fractions were isolated and 80 μg per sample were run on an SDS-PAGE gel and imaged for Bocillin-FL (top). After imaging the gels were Coomassie stained for total protein quantification (bottom). See Figure S1 for quantification.

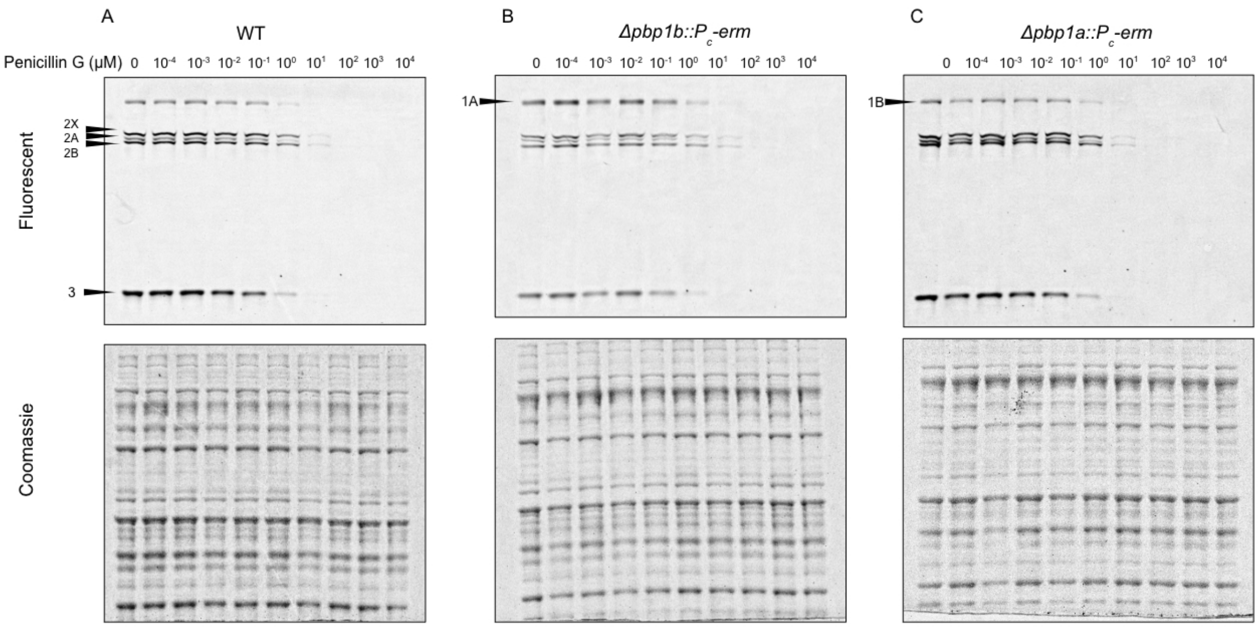


**Figure S5**. Bocillin End Point Assay representative gel images for penicillin G inhibition. Live *S. pneumoniae* WT, Δ*pbp1B* (B), or *Δpbp1A* (C), cells were treated with Bocillin-FL (600 nM) and a concentration gradient of penicillin G ($10^{-4}$-$10^{4}$ μM). After incubation, the cells were lysed with lysozyme and sonication. The membrane fractions were isolated and 80 μg per sample were run on an SDS-PAGE gel and imaged for Bocillin-FL (top). After imaging the gels were Coomassie stained for total protein quantification (bottom). See Figure S1 for quantification.

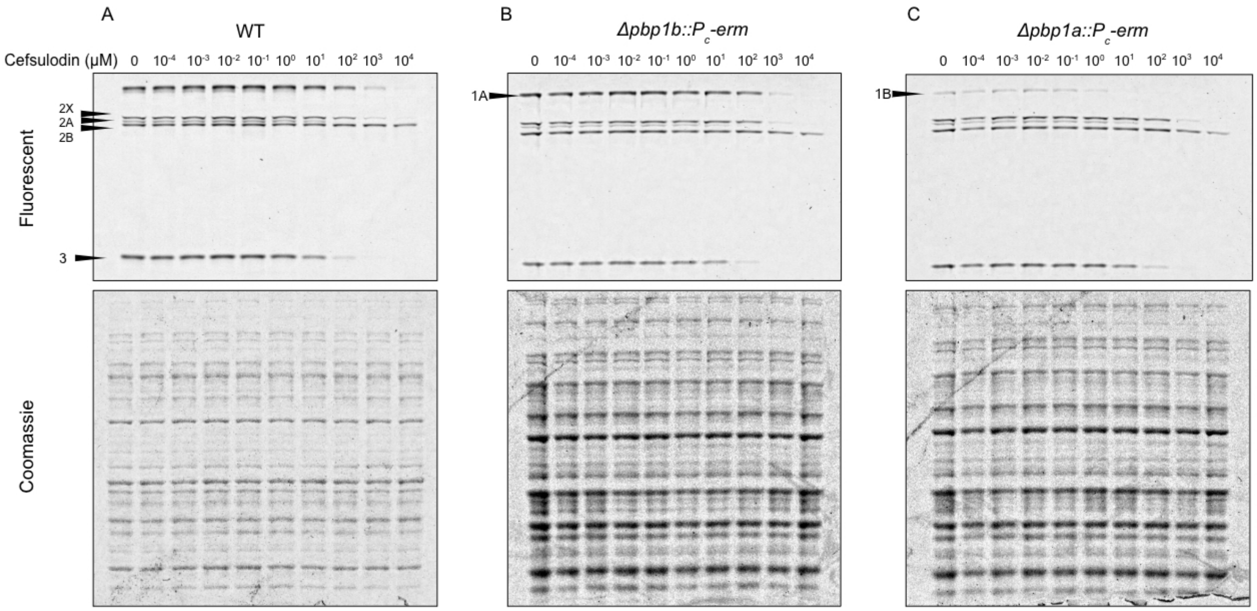


**Figure S6**. Bocillin End Point Assay representative gel images for cefsulodin inhibition. Live *S. pneumoniae* WT, Δ*pbp1B* (B), or *Δpbp1A* (C), cells were treated with Bocillin-FL (600 nM) and a concentration gradient of cefsulodin ($10^{-4}$-$10^{4}$ μM). After incubation, the cells were lysed with lysozyme and sonication. The membrane fractions were isolated and 80 μg per sample were run on an SDS-PAGE gel and imaged for Bocillin-FL (top). After imaging the gels were Coomassie stained for total protein quantification (bottom). See Figure S1 for quantification.

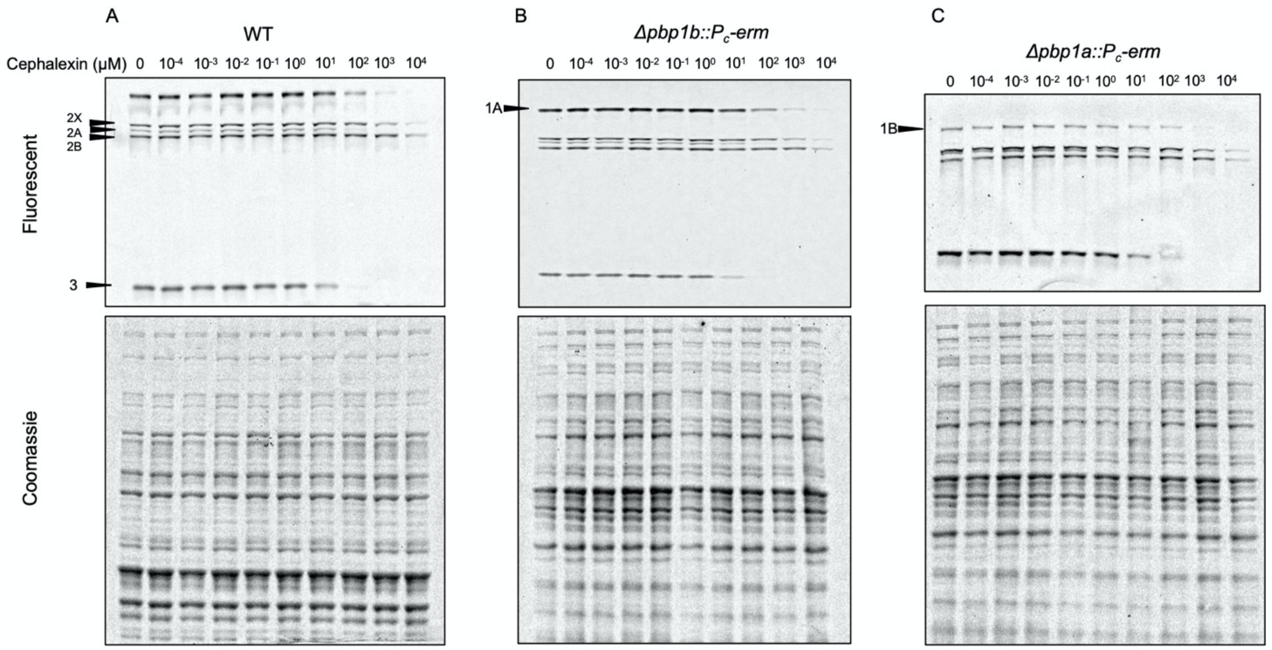


**Figure S7**. Bocillin End Point Assay representative gel images for cephalexin inhibition. Live *S. pneumoniae* WT, Δ*pbp1B* (B), or *Δpbp1A* (C), cells were treated with Bocillin-FL (600 nM) and a concentration gradient of cephalexin ($10^{-4}$-$10^{4}$ μM). After incubation, the cells were lysed with lysozyme and sonication. The membrane fractions were isolated and 80 μg per sample were run on an SDS-PAGE gel and imaged for Bocillin-FL (top). After imaging the gels were Coomassie stained for total protein quantification (bottom). See Figure S1 for quantification.

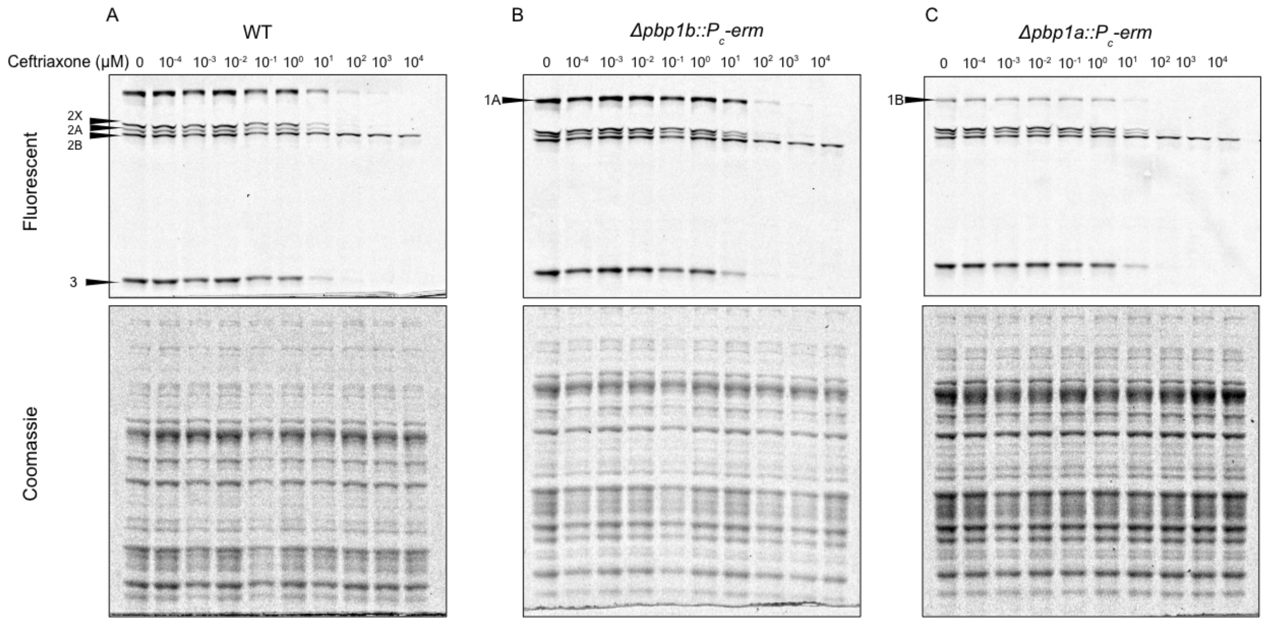


**Figure S8**. Bocillin End Point Assay representative gel images for ceftriaxone inhibition. Live *S. pneumoniae* WT, Δ*pbp1B* (B), or *Δpbp1A* (C), cells were treated with Bocillin-FL (600 nM) and a concentration gradient of ceftriaxone ($10^{-4}$-$10^{4}$ μM). After incubation, the cells were lysed with lysozyme and sonication. The membrane fractions were isolated and 80 μg per sample were run on an SDS-PAGE gel and imaged for Bocillin-FL (top). After imaging the gels were Coomassie stained for total protein quantification (bottom). See Figure S1 for quantification.

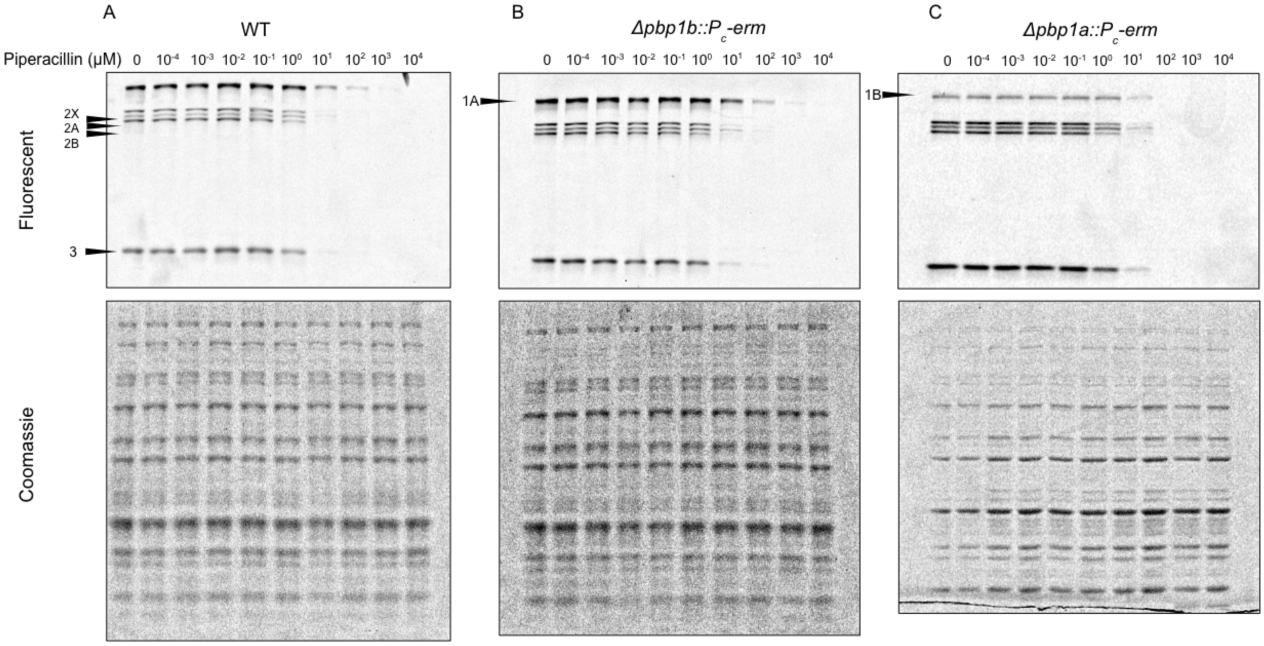


**Figure S9**. Bocillin End Point Assay representative gel images for piperacillin inhibition. Live *S. pneumoniae* WT, Δ*pbp1B* (B), or *Δpbp1A* (C), cells were treated with Bocillin-FL (600 nM) and a concentration gradient of piperacillin ($10^{-4}$-$10^{4}$ μM). After incubation, the cells were lysed with lysozyme and sonication. The membrane fractions were isolated and 80 μg per sample were run on an SDS-PAGE gel and imaged for Bocillin-FL (top). After imaging the gels were Coomassie stained for total protein quantification (bottom). See Figure S1 for quantification.

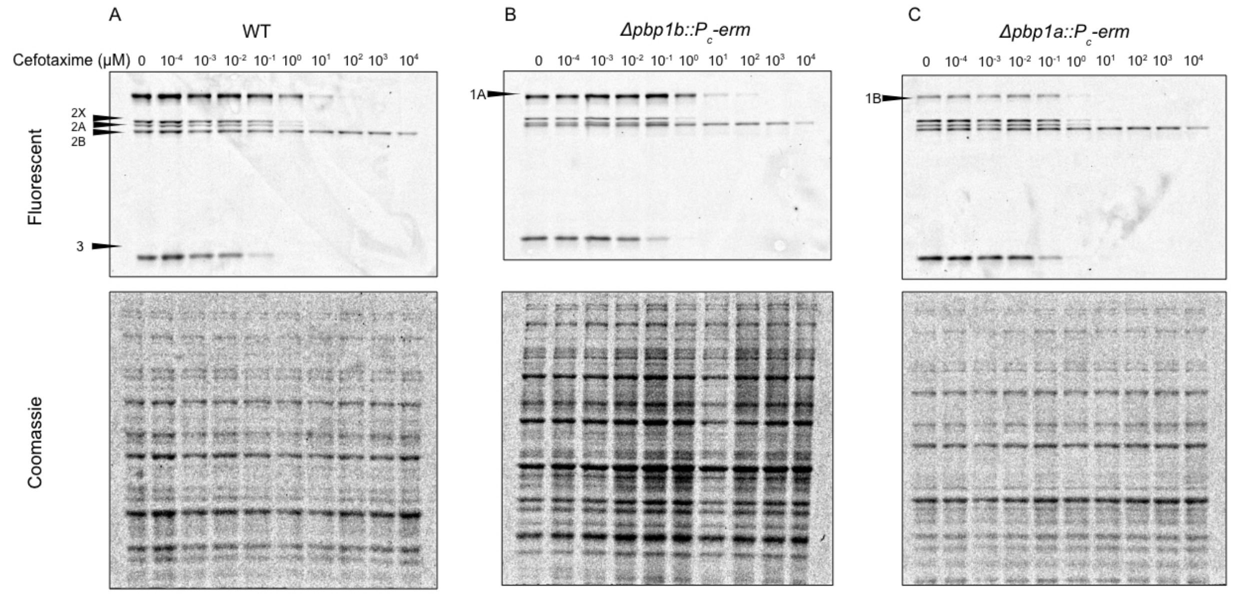


**Figure S10**. Bocillin End Point Assay representative gel images for cefotaxime inhibition. Live *S. pneumoniae* WT, Δ*pbp1B* (B), or *Δpbp1A* (C), cells were treated with Bocillin-FL (600 nM) and a concentration gradient of cefotaxime ($10^{-4}$-$10^{4}$ μM). After incubation, the cells were lysed with lysozyme and sonication. The membrane fractions were isolated and 80 μg per sample were run on an SDS-PAGE gel and imaged for Bocillin-FL (top). After imaging the gels were Coomassie stained for total protein quantification (bottom). See Figure S1 for quantification.

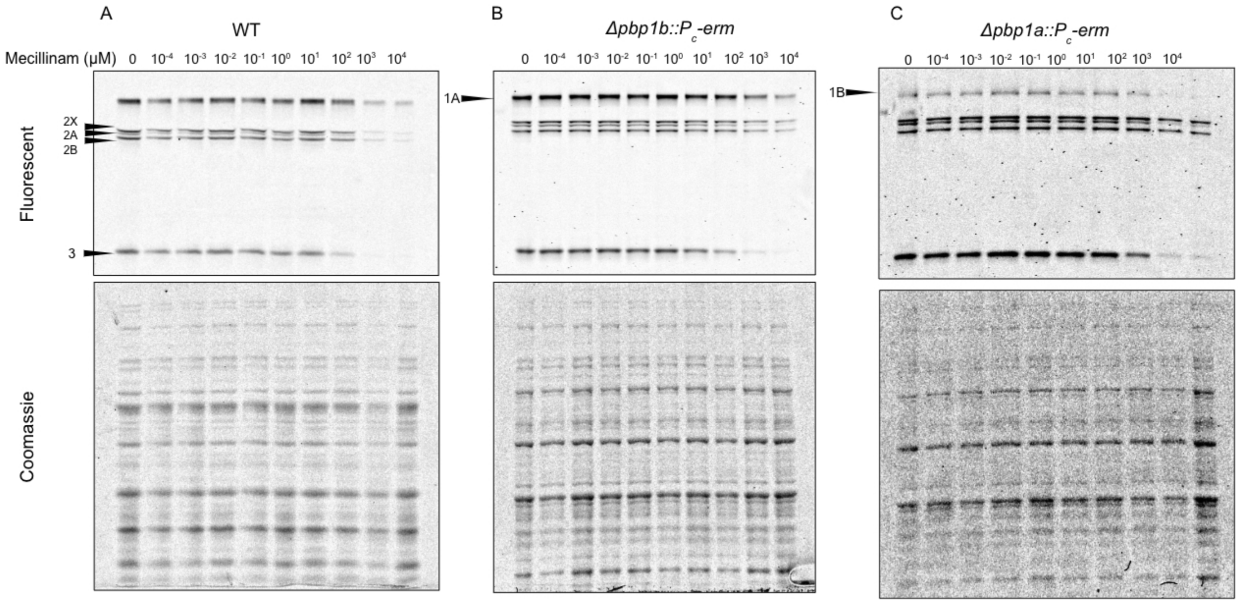


**Figure S11**. Bocillin End Point Assay representative gel images for mecillinam inhibition. Live *S. pneumoniae* WT, Δ*pbp1B* (B), or *Δpbp1A* (C), cells were treated with Bocillin-FL (600 nM) and a concentration gradient of mecillinam ($10^{-4}$-$10^{4}$ µM). After incubation, the cells were lysed with lysozyme and sonication. The membrane fractions were isolated and 80 µg per sample were run on an SDS-PAGE gel and imaged for Bocillin-FL (top). After imaging the gels were Coomassie stained for total protein quantification (bottom). See Figure S1 for quantification.
